# Learning Sparse Log-Ratios for High-Throughput Sequencing Data

**DOI:** 10.1101/2021.02.11.430695

**Authors:** Elliott Gordon-Rodriguez, Thomas P. Quinn, John P. Cunningham

## Abstract

The automatic discovery of sparse biomarkers that are associated with an outcome of interest is a central goal of bioinformatics. In the context of high-throughput sequencing (HTS) data, and *compositional data* (CoDa) more generally, an important class of biomarkers are the log-ratios between the input variables. However, identifying predictive log-ratio biomarkers from HTS data is a combinatorial optimization problem, which is computationally challenging. Existing methods are slow to run and scale poorly with the dimension of the input, which has limited their application to low- and moderate-dimensional metagenomic datasets. Building on recent advances from the field of deep learning, we present *CoDaCoRe*, a novel learning algorithm that identifies sparse, interpretable, and predictive log-ratio biomarkers. Our algorithm exploits a *continuous relaxation* to approximate the underlying combinatorial optimization problem. This relaxation can then be optimized efficiently using the modern ML toolbox, in particular, gradient descent. As a result, CoDaCoRe runs several orders of magnitude faster than competing methods, all while achieving state-of-the-art performance in terms of predictive accuracy and sparsity. We verify the outperformance of CoDaCoRe across a wide range of microbiome, metabolite, and microRNA benchmark datasets, as well as a particularly high-dimensional dataset that is outright computationally intractable for existing sparse log-ratio selection methods.^1^

## 1 Introduction

High-throughput sequencing (HTS) technologies have enabled the relative quantification of the different bacteria, metabolites, or genes, that are present in a biological sample. However, the nature of these recording technologies results in *sequencing biases* that complicate the analysis of HTS data. In particular, HTS data come as counts, whose totals are constrained to the capacity of the measuring device. These totals are an artifact of the measurement process, and do not depend on the subject being measured. Hence, HTS counts arguably should be interpreted in terms of *relative abundance*; in statistical terminology, it follows that HTS data are an instance of *compositional data* (CoDa) (Gloor *et al.*, 2016, 2017; Quinn *et al.*, 2018, 2019; Calle, 2019).

Mathematically, CoDa can be defined as a set of non-negative vectors whose totals are uninformative. Since the seminal work of Aitchison (1982), the statistical analysis of CoDa has become a discipline in its own right (Pawlowsky-Glahn and Egozcue, 2006; Pawlowsky-Glahn and Buccianti, 2011; Pawlowsky-Glahn *et al.*, 2015). But why does CoDa deserve special treatment? Unlike unconstrained real-valued data, the compositional nature of CoDa results in each variable becoming negatively correlated to all others (increasing one component of a composition implies a relative decrease of the other components). It is well known that, as a result, the usual measures of association and feature attribution are problematic when applied to CoDa (Pearson, 1896; Filzmoser *et al.*, 2009; Van den Boogaart and Tolosana-Delgado, 2013; Lovell *et al.*, 2015). Consequently, bespoke methods are necessary for a valid statistical analysis (Gloor *et al.*, 2017). Indeed, the application of CoDa methodology to HTS data, especially microbiome data, has become increasingly popular in recent years (Fernandes *et al.*, 2013, 2014; Rivera-Pinto *et al.*, 2018; Quinn *et al.*, 2019; Calle, 2019; Quinn *et al.*, 2021).

The standard approach for analyzing CoDa is based on applying *log-ratio* transformations to map our data onto unconstrained Euclidean space, where the usual tools of statistical learning apply (Pawlowsky-Glahn and Egozcue, 2006). The choice of the log-ratio transform offers the necessary property of scale invariance, but in the CoDa literature it holds primacy for a variety of other technical reasons, including *subcompositional coherence* (Aitchison, 1982; Pawlowsky-Glahn and Buccianti, 2011; Egozcue and Pawlowsky-Glahn, 2019). Logratios can be taken over pairs of input variables (Aitchison, 1982; Greenacre, 2019b; Bates and Tibshirani, 2019) or aggregations thereof, typically geometric means (Aitchison, 1982; Egozcue *et al.*, 2003; Egozcue and Pawlowsky-Glahn, 2005; Rivera-Pinto *et al.*, 2018; Quinn and Erb, 2019) or summations (Greenacre, 2019a, 2020; Quinn and Erb, 2020). The resulting features work well empirically, but also imply a clear interpretation: a log-ratio is a single composite score that expresses the overall quantity of one sub-population as compared with another. For example, in microbiome HTS data, the relative weights between sub-populations of related microorganisms are commonly used as clinical biomarkers (Rahat-Rozenbloom *et al.*, 2014; Crovesy *et al.*, 2020; Magne *et al.*, 2020). When the log-ratios are sparse, meaning they are taken over a small number of input variables, they define biomarkers that are particularly intuitive to understand, a key desiderata for predictive models that are of clinical relevance (Goodman and Flaxman, 2017).

### Thus, learning sparse log-ratios is a central problem in CoDa

This problem is especially challenging in the context of HTS data, due to its high dimensionality (ranging from 100 to over 10,000 variables). Existing methods rely on stepwise search (Rivera-Pinto *et al.*, 2018; Greenacre, 2019b) or evolutionary algorithms (Quinn and Erb, 2020; Prifti *et al.*, 2020), which scale poorly with the dimension of the input. These algorithms are prohibitively slow for most HTS datasets, and thus there is a new demand for sparse and interpretable models that scale to high dimensions (Li, 2015; Cammarota *et al.*, 2020; Susin *et al.*, 2020).

This demand motivates the present work, in which we present CoDaCoRe, a novel learning algorithm for **Co**mpositional **Da**ta via **Co**ntinuous **Re**laxations. CoDaCoRe builds on recent advances from the deep learning literature on *continuous relaxations* of discrete latent variables (Jang *et al.*, 2016; Maddison *et al.*, 2017; Linderman *et al.*, 2018; Mena *et al.*, 2018; Potapczynski *et al.*, 2020); we design a novel relaxation that approximates a combinatorial optimization problem over the set of log-ratios. In turn, this approximation can be optimized efficiently using gradient descent, and subsequently discretized to produce a sparse log-ratio biomarker, thus dramatically reducing runtime without sacrificing interpretability nor predictive accuracy. The main contributions of our method can be summarized as follows:

- **Computational efficiency.** CoDaCoRe scales linearly with the dimension of the input. It runs several orders of magnitude faster than its competitors.
- **Interpretability.** CoDaCoRe identifies a set of log-ratios that are sparse, biologically meaningful, and ranked in order of importance. Our model is highly interpretable, and much sparser, relative to competing methods of similar accuracy and computational complexity.
- **Predictive accuracy.** CoDaCoRe achieves better out-of-sample accuracy than existing CoDa methods, and performs similarly to state-of-the-art black-box classifiers (which are neither sparse nor interpretable).
- **Ease of use.** We devise an adaptive learning rate scheme that enables CoDaCoRe to converge reliably, requiring no additional hyperparameter tuning. As a result, deploying CoDaCoRe on novel datasets is straightforward:

~~~
devtools::install_github(“egr95/R-codacore”)
model <- codacore(x, y)
print(model)
plot(model)
~~~

## 2 Background

Our work focuses on the supervised learning problem **x**_*i*_ ↦ *y_i_*, where the inputs **x**_*i*_ are HTS data (or any CoDa), and the outputs *y_i_* are the outcome of interest. For many microbiome applications, **x**_*i*_ represents a vector of frequencies of the different species of bacteria that compose the microbiome of the *i*th subject. In other words, *x_ij_* denotes the abundance of the *j*th species (of which there are *p* total) in the *i*th subject. The response *y_i_* is often a binary variable indicating whether the *i*th subject belongs to the case or the control groups (e.g., sick vs. healthy). For HTS data, the input frequencies *x_ij_* arise from an inexhaustive sampling procedure, so that the totals 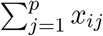 are arbitrary and the components should only be interpreted in relative terms (i.e., as CoDa) (Gloor and Reid, 2016; Gloor *et al.*, 2017; Quinn *et al.*, 2018; Calle, 2019). While many of our applications pertain to microbiome data, our method applies more generally to any high-dimensional HTS data, including those produced by *Liquid Chromatography Mass Spectrometry* (Filzmoser and Walczak, 2014).

### 2.1 Log-Ratio Analysis

Our goal is to obtain sparse log-ratio transformed features that can be passed to a downstream classifier or regression function. As discussed, these log-ratios will result in interpretable features and scale-invariant models (that are also subcompositionally coherent), thus satisfying the key requirements for valid statistical inference in the context of CoDa. The simplest such choice is the *pairwise log-ratio*, defined as log(*x_ij_*+ /*x_ij–_*), where *j*^+^ and *j*^−^ denote the indexes of a pair of input variables (Aitchison, 1982). Note that the ratio cancels out any scaling factor applied to **x**_*i*_, preserving only the relative information, while the log transformation ensures the output is (unconstrained) real-valued. There are many such (*j*^+^, *j*^−^) pairs (to be precise, *p*(*p* – 1)/2 = *O*(*p*^2^) of them). In order to select good pairwise log-ratios from a set of input variables, Greenacre (2019b) proposed a greedy step-wise search algorithm. This method produces a sparse and interpretable set of features, but it is prohibitively slow on high-dimensional datasets, as a result of the step-wise algorithm scaling quadratically in the dimension of the input. A heuristic search algorithm that is less accurate but computationally faster has been developed as part of Quinn *et al.* (2017), though its computational cost is still troublesome (as we shall see in Section 4).

#### 2.1.1 Balances

Recently, a class of log-ratios known as *balances* (Egozcue and Pawlowsky-Glahn, 2005) have become of interest in microbiome applications, due to their interpretability as the relative weight between two sub-populations of bacteria (Morton *et al.*, 2019a; Quinn and Erb, 2019). Balances are defined as the log-ratios between geometric means of two subsets of the input variables:^2^

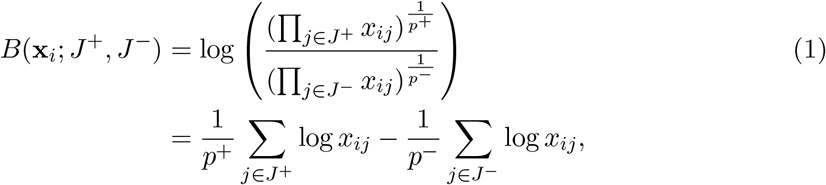

where *J*^+^ and *J*^−^ denote a pair of disjoint subsets of the indices {1,...,*p*}, and *p*^+^ and *p*^−^ denote their respective sizes. For example, in microbiome data, *J*^+^ and *J*^−^ are groups of bacteria species that may be related by their environmental niche (Morton *et al.*, 2017) or genetic similarity (Silverman *et al.*, 2017; Washburne *et al.*, 2017). Note that when *p*^+^ = *p*^−^ =1 (i.e., *J*^+^ and *J*^−^ each contain a single element), *B*(**x**; *J*^+^, *J*^−^) reduces to a pairwise log-ratio. By allowing for the aggregation of more than one variable in the numerator and denominator of the log-ratio, balances provide a far richer set of features that allows for more flexible models than pairwise log-ratios. Insofar as the balances are taken over a small number of variables (i.e., *J*^+^ and *J^−^* are sparse), they also provide highly interpretable biomarkers.

The *selbal* algorithm (Rivera-Pinto *et al.*, 2018) has gained popularity as a method for automatically identifying balances that predict a response variable. However, this algorithm is also based on a greedy step-wise search through the combinatorial space of subset pairs (*J*^+^, *J*^−^), which scales poorly in the dimension of the input and becomes prohibitively slow for many HTS datasets (Susin *et al.*, 2020).

#### 2.1.2 Amalgamations

An alternative to balances, known as *amalgamation*, is defined by aggregating components through summation:

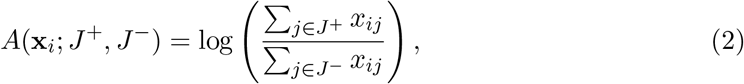

where again *J*^+^ and *J*^−^ denote disjoint subsets of the input components. Amalgamations have the advantage of reducing the dimensionality of the data through an operation, the sum, that some authors argue is more interpretable than a geometric mean (Greenacre, 2019a; Greenacre *et al.*, 2020). On the other hand, amalgamations can be less effective than balances for identifying components that are statistically important, but small in magnitude, e.g., rare bacteria species (since small terms will have less impact on a summation than on a product).

Recently, Greenacre (2020) has advocated for the use of expert-driven amalgamations, using domain knowledge to construct the relevant features. On the other hand, Quinn and Erb (2020) proposed *amalgam*, an evolutionary algorithm to automatically identify amalgamated log-ratios (Eq. 2) that are predictive of a response variable. However, this algorithm does not scale to high-dimensional data (albeit, comparing favorably to selbal), nor does it produce sparse models (hindering interpretability of the results). A similar evolutionary algorithm can be found in Prifti *et al.* (2020), however their model is not scale invariant, as is required by most authors in the field (Pawlowsky-Glahn and Egozcue, 2006).

#### 2.1.3 Other Related Work

CoDa methodology has also recently attracted interest from the machine learning community (Tolosana-Delgado *et al.*, 2019; Quinn *et al.*, 2020; Gordon-Rodriguez *et al.*, 2020a,b; Templ, 2020). Relevant to us is *DeepCoDA* (Quinn *et al.*, 2020), which combines self-explaining neural networks with log-ratio transformed features. In particular, DeepCoDA learns a set of *log-contrasts*, in which the numerator and denominator are defined as *unequally weighted* geometric averages of components Egozcue and Pawlowsky-Glahn (2016). As a result of this weighting, DeepCoDA loses much of the interpretability and intuitive appeal of balances (or amalgamations), which is exacerbated by its lack of sparsity. Moreover, like most deep architectures, DeepCoDA is sensitive to initialization and optimization hyperparameters (which limits its ease of use) and is susceptible to overfitting (which can further compromise interpretability of the model).

The special case of a linear log-contrast model has been referred to as *Coda-lasso*, and was separately proposed by Lu *et al.* (2019). While Coda-lasso scales better than selbal, it has been found to perform worse in terms of predictive accuracy (Susin *et al.*, 2020). More importantly, Coda-lasso is still prohibitively slow on the high-dimensional HTS data that we wish to consider. Last, we highlight another common set of features that are also a special case of log-contrasts: *centered-log-ratios* (CLR), where each input variable is divided by the geometric mean of *all* input variables (Aitchison, 1982). Models using these features, such as CLR-lasso (Susin *et al.*, 2020), can be accurate and computationally efficient, however they are inherently not sparse and are difficult to interpret scientifically (Greenacre, 2019a).

## 3 Methods

We now present CoDaCoRe, a novel learning algorithm for HTS data, and more generally, high-dimensional CoDa. Unlike existing methods, CoDaCoRe is simultaneously scalable, interpretable, sparse, and accurate. We compare the relative merits of CoDaCoRe and its competitors in Table 1.

**Table 1:** Qualitative comparison of learning algorithms, ordered from most sparse (top) to least (bottom). CoDaCoRe is the only method that performs on all of our criteria. See Table 2 for a corresponding quantitative comparison.

|  | Scalability | Interpretability | Sparsity | Accuracy |
| --- | --- | --- | --- | --- |
| **CoDaCoRe (ours)** | + | + | + | + |
| Pairwise log-ratios (Greenacre, 2019b) | − | + | + | − |
| Selbal (Rivera-Pinto *et al.*, 2018) | − | + | + | · |
| Lasso | + | · | + | − |
| Coda-lasso (Lu *et al.*, 2019) | − | · | · | · |
| Amalgam (Quinn and Erb, 2020) | − | + | − | · |
| DeepCoDA (Quinn *et al.*, 2020) | · | · | − | · |
| CLR-lasso (Susin *et al.*, 2020) | + | − | − | + |
| Black-box (e.g., Random Forest, XGB) | + | − | − | + |

### 3.1 Optimization Problem

In its basic formulation, CoDaCoRe learns a regression function of the form:

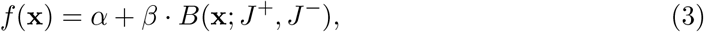

where *B* denotes a balance (Eq. 1), and *α* and *β* are scalar parameters. This regression function can be thought of in two stages: first we take the input and use it to compute a balance score, second we feed the balance score to a logistic regression classifier. For clarity, we will restrict our exposition to this formulation, but note that our algorithm can be applied equally to learn amalgamations instead of balances (see Section 3.6), as well as generalizing straightforwardly to nonlinear functions (provided they are suitably parameterized and differentiable).

Let *L*(*y, f*) denote the cross-entropy loss, with *f* ∈ ℝ given in logit space, or in other words:

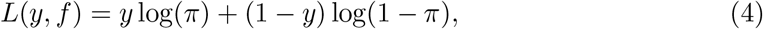

where:

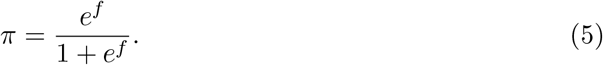

The goal of CoDaCoRe is to find the balance that is maximally associated with the response. Mathematically, this can be written as:

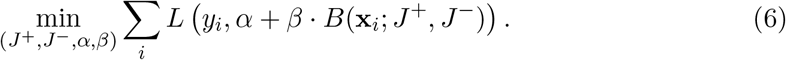

This objective function may look similar to a univariate logistic regression, however our problem is complicated by the joint optimization over the subsets *J*^+^ and *J*^−^, which determine the input variables that compose the balance. Note that the number of possible subsets of *p* variables is 2^*p*^, so the set of possible balances is greater than 2^*p*^ and grows *exponentially* in *p*. Exact optimization is therefore computationally intractable for any but the smallest of datasets, and an approximate solution is required. Selbal corresponds to one such approximation, offering *quadratic* complexity in *p*, which is practical for low- to moderate-dimensional datasets (*p* < 100), but does not scale to high dimensions (*p* > 1,000). As we shall now see, CoDaCoRe represents a critical improvement, achieving *linear* complexity in *p* which dramatically reduces runtime and enables, for the first time, the use of balances and amalgamations for the analysis of high-dimensional HTS data.

### 3.2 Continuous Relaxation

*The key insight of CoDaCoRe is to approximate our combinatorial optimization problem (Eq. 6) with a continuous relaxation that can be trained efficiently by gradient descent.* Our relaxation is inspired by recent advances in deep learning models with discrete latent variables (Jang *et al.*, 2016; Maddison *et al.*, 2017; Linderman *et al.*, 2018; Mena *et al.*, 2018; Potapczynski *et al.*, 2020). However, we are not aware of any similar proposals for optimizing over disjoint subsets, nor for learning balances or amalgamations in the context of CoDa.

Our relaxation is parameterized by an unconstrained vector of “assignment weights”, **w** ∈ ℝ^*p*^, with one scalar parameter per input dimension (e.g., one weight per bacteria species). The weights are mapped to a vector of “soft assignments” via:

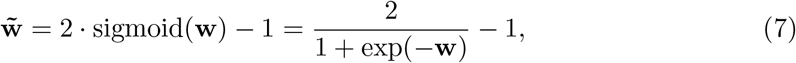

where the sigmoid is applied component-wise. Intuitively, large positive weights will max out the sigmoid, leading to soft assignments close to +1, whereas large negative weights will zero out the sigmoid, resulting in soft assignments close to –1. This mapping is akin to softly assigning input variables to the groups *J*^+^ and *J*^−^, respectively.

Let us write 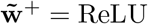 and 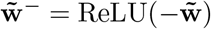 for the (component-wise) positive and negative parts of 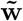, respectively. We approximate balances (Eq. 1) with the following relaxation:

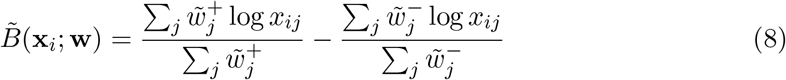

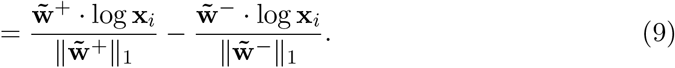

In other words, we approximate the geometric averages over subsets of the inputs, by *weighted* geometric averages over all components (compare Equations 1 and 8).

Crucially, this relaxation is differentiable in **w**, allowing us to construct a surrogate objective function that can be optimized jointly in (**w**, *α, β*) by gradient descent:

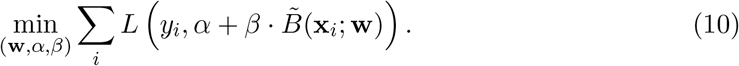

Moreover, the computational cost of differentiating this objective function scales linearly in the dimension of **w**, which overall results in linear scaling for our algorithm. We also note that the functional form of our relaxation (Eq. 8) can be exploited in order to select the learning rate adaptively (i.e., without tuning), resulting in robust convergence across all real and simulated datasets that we considered. We defer the details of our implementation of gradient descent to the Supplement (Section A).

### 3.3 Discretization

While a set of features in the form of Eq. 8 may perform accurate classification, a weighted geometric average over all input variables is much harder for a biologist to interpret (and less intuitively appealing) than a bona fide balance over a small number of variables. For this reason, CoDaCoRe implements a “discretization” procedure that exploits the information learned by the soft assignment vector 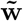, in order to efficiently identify a pair of sparse subsets, 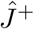 and 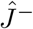, which will define a balance.

The most straightforward way to convert the (soft) assignment 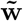 into a (hard) pair of subsets is by fixing a threshold *t* ∈ (0,1):

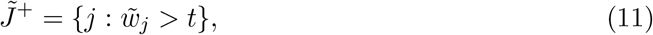

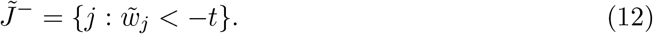

Note that given a trained 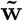 and a fixed threshold *t*, we can evaluate the quality of the corresponding balance 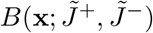 (resp. amalgamation) by optimizing Eq. 6 over (*α, β*) alone, i.e., fitting a linear model. Computationally, fitting a linear model is much faster than optimizing Eq. 10, and can be done repeatedly for a range of values of *t* with little overhead. In CoDaCoRe, we combine this strategy with cross-validation in order to select the threshold, 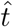, that optimizes predictive performance (see Section A of the Supplement for full detail). Finally, the trained regression function is:

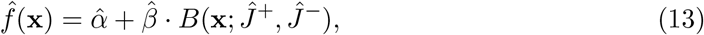

where 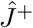 and 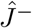 are the subsets corresponding to the optimal threshold 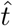, and 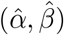 are the coefficients obtained by regressing *y_i_* against 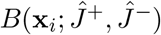 on the entire training set.

### 3.4 Regularization

Note from Equations 11 and 12 that larger values of *t* result in fewer input variables assigned to the balance 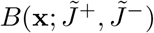, i.e., a sparser model. Thus, CoDaCoRe can be regularized simply by making 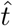 larger. Similar to lasso regression, CoDaCoRe uses the *1-standard-error* rule: namely, to pick the sparsest model (i.e., the highest *t*) with mean cross-validated score within 1 standard error of the optimum (Friedman *et al.*, 2001). Trivially, this rule can be generalized to a λ-standard-error rule (to pick the sparsest model within λ standard errors of the optimum), where λ becomes a regularization hyperparameter that can be tuned by the practitioner if so desired (with lower values trading off some sparsity in exchange for predictive accuracy). In our public implementation, λ = 1 is our default value, and this is used throughout our experiments (except where we indicate otherwise). In practice, lower values (e.g., λ = 0) can be useful when the emphasis is on predictive accuracy rather than interpretability or sparsity, though our benchmarks showed competitive performance for any λ ∈ [0,1].

**Algorithm 1.**
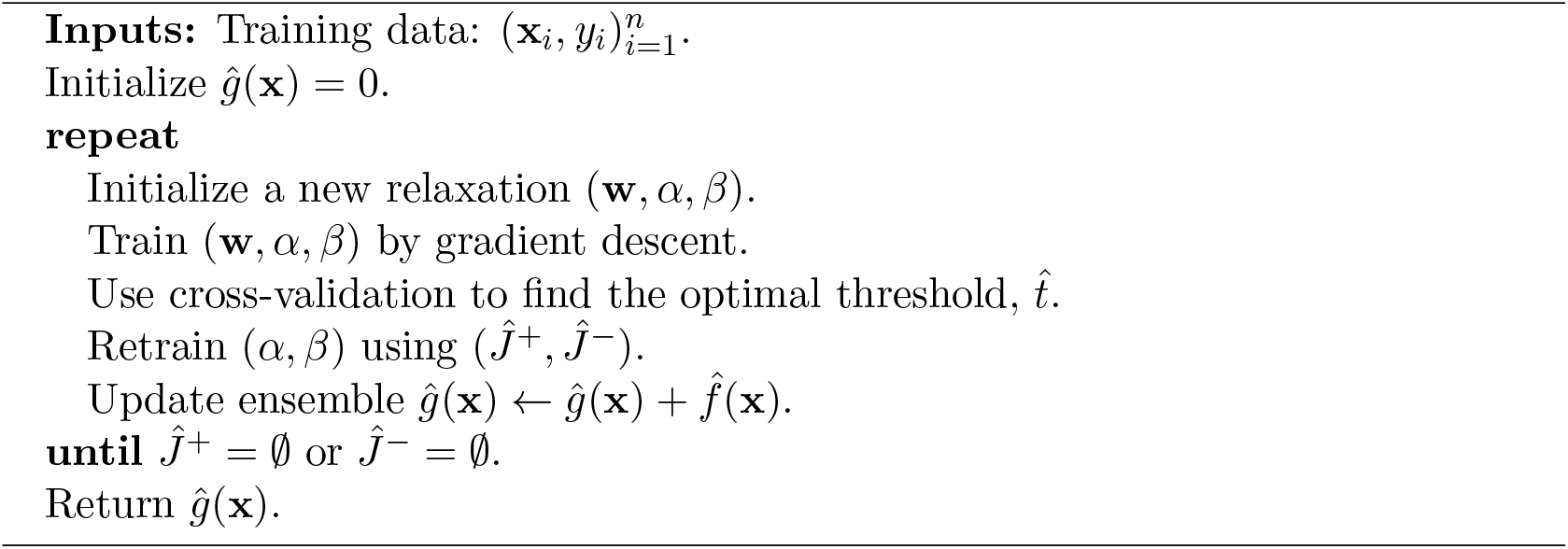
CoDaCoRe

### 3.5 CoDaCoRe Algorithm

The computational efficiency of our continuous relaxation allows us to train multiple regressors of the form of Eq. 13 within a single model. In the full CoDaCoRe algorithm, we ensemble multiple such regressors in a stage-wise additive fashion, where each successive balance is fitted on the residual from the current model. Thus, CoDaCoRe identifies a *sequence* of balances, in decreasing order of importance, each of which is sparse and interpretable. Training terminates when an additional relaxation (Eq. 8) cannot improve the cross-validation score relative to the existing ensemble (equivalently, when we obtain 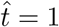). Typically, only a small number of balances is required to capture the signal in the data, and as a result CoDaCoRe produces very sparse models overall, further enhancing interpretability. Our procedure is summarized in Algorithm 1.

### 3.6 Amalgamations

CoDaCoRe can be used to learn amalgamations (Eq. 2) much in the same way as for balances (the choice of which to use depending on the goals of the biologist). In this case, our relaxation is defined as:

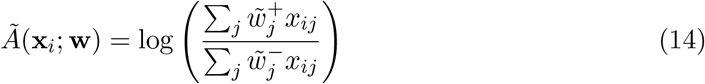

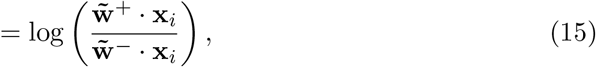

i.e., we approximate summations over subsets of the inputs, with *weighted* summations over all components (compare Eq. 2 and Eq. 14). The rest of the argument follows verbatim, replacing *B*(·) with *A*(·) and *B*(·) with *A*(·) in Equations 3, 6, 10, and 13.

### 3.7 Extensions

Our model allows for a number of extensions:

- *Unsupervised learning.* By means of a suitable unsupervised loss function, CoDaCoRe can be extended to unlabelled datasets, 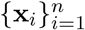, as a method for identifying log-ratios that provide a useful low-dimensional representation. Such a method would automatically provide a scalable alternative to several existing dimensionality reduction techniques for CoDa (Pawlowsky-Glahn *et al.*, 2011; Mert *et al.*, 2015; Martín-Fernandez *et al.*, 2018; Martino *et al.*, 2019; Quinn *et al.*, 2021).
- *Incorporating confounders.* In addition to 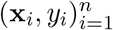, in some applications the effect of additional (non-compositional) predictors, **z**_*i*_, is also of interest. In this case, the effect of **z**_*i*_ can be “partialled out” a priori by first regressing *y_i_* on **z**_*i*_ alone, and using this regression as the initialization of the CoDaCoRe ensemble. Alternatively, **z**_*i*_ can also be modeled jointly in Equations 3 and 13 (e.g., by adding a linear term *γ* · **z**_*i*_) (Forslund *et al.*, 2015; Noguera-Julian *et al.*, 2016; Rivera-Pinto *et al.*, 2018).
- *Nonlinear regression functions.* Our method extends naturally to nonlinear regression functions of the form *f*(**x**) = *h_θ_*(*B*(**x**; *J*^+^, *J*^−^)), where *h_θ_* is a parameterized differentiable family, including neural networks (Morton *et al.*, 2019b; Quinn *et al.*, 2020).
- *Applications to non-compositional data.* Aggregations of parts can be useful outside the realm of CoDa; for example, an amalgamation applied to a categorical variable with many levels represents a grouping of the categories (Bondell and Reich, 2009; Gertheiss and Tutz, 2010; Tutz and Gertheiss, 2016).

## 4 Experiments

We evaluate CoDaCoRe on a collection of 25 benchmark datasets including 13 datasets from the *Microbiome Learning Repo* (Vangay *et al.*, 2019), and 12 microbiome, metabolite, and microRNA datasets curated by Quinn and Erb (2019). These data vary in dimension from 48 to 3,090 input variables (see Section B of the Supplement for a full description). For each dataset, we fit CoDaCoRe on 20 random 80/20 train/test splits, sampled with stratification by case-control (He and Ma, 2013). We compare against:

- Interpretable models (Sections 2.1.1 and 2.1.2): pairwise log-ratios (Greenacre, 2019b)^3^, selbal (Rivera-Pinto *et al.*, 2018), and amalgam (Quinn and Erb, 2020). We also consider lasso logistic regression (with regularization parameter chosen by crossvalidation with the 1-standard-error rule).
- Other CoDa models (Section 2.1.3): Coda-lasso (Lu *et al.*, 2019), DeepCoDA (Quinn *et al.*, 2020), and CLR-lasso (Susin *et al.*, 2020). Note that these methods learn weighted geometric averages over a large number of input variables, which are evidently not as straightforward to interpret as simple balances or amalgamations.
- Black box classifiers: Random Forest and XGBoost (Chen and Guestrin, 2016), where we tune the model complexity parameters by cross-validation (subsample size and early stopping, respectively).

### 4.1 Results

We evaluate the quality of our models across the following criteria: computational efficiency (as measured by runtime), sparsity (as measured by the percentage of input variables that are active in the model), and predictive accuracy (as measured by out-of-sample accuracy, ROC AUC, and F1 score). Table 2 provides an aggregated summary of the results; CoDaCoRe (with balances) is performant on all metrics. Indeed, our method provides the only interpretable model that is simultaneously scalable, sparse, and accurate. Detailed performance metrics on each of the 25 datasets are provided in Section C of the Supplement.

**Table 2:** Evaluation metrics shown for each method, averaged over 25 datasets × 20 random train/test splits. Standard errors are computed independently on each dataset, and then averaged over the 25 datasets. The models are ordered by sparsity, i.e., percentage of active input variables. CoDaCoRe (with balances) is the only learning algorithm that is simultaneously fast, sparse, and accurate. The bottom rows show best-in-class black-box classifiers, which can be thought of as providing an approximate upper bound on the predictive accuracy of any interpretable model.

|  | Runtime (s) | Active Vars (%) | Accuracy (%) | AUC (%) | F1 (%) |
| --- | --- | --- | --- | --- | --- |
| Majority class | 0.0±0.0 | 0.0±0.0 | 62.5±0.0 | 50.0±0.0 | · |
| CoDaCoRe - Balances (ours) | **4.5±0.4** | **1.9±0.3** | 75.2±2.4 | 79.5±2.6 | 73.7±2.6 |
| CoDaCoRe - Amalgamations (ours) | **4.4±0.4** | 1.9±0.3 | 71.8±2.4 | 74.5±2.8 | 69.8±2.9 |
| selbal (Rivera-Pinto *et al.*, 2018) | 79,033.7±2,094.1 | 2.4±0.2 | 61.2±1.9 | 80.0±2.4 | 70.9±1.1 |
| Pairwise Log-ratios (Greenacre, 2019b) | 14,207.0±1,038.4 | 2.5±0.4 | 73.3±1.7 | 75.2±2.4 | 67.8±3.0 |
| Lasso | **1.6±0.1** | 4.4±0.6 | 72.4±1.7 | 75.2±2.3 | 65.2±3.7 |
| CoDaCoRe - Balances with $\lambda = 0$ (ours) | **9.8±2.2** | 6.1±0.7 | **77.6±2.2** | **82.0±2.3** | **76.0±2.5** |
| Coda-lasso (Lu *et al.*, 2019) | 1,043.0±55.4 | 19.7±2.7 | 72.5±2.3 | 78.0±2.4 | 64.2±4.4 |
| amalgam (Quinn and Erb, 2020) | 7,360.5±209.8 | 87.6±2.1 | 74.4±2.5 | 78.2±2.7 | 73.9±2.8 |
| DeepCoDA (Quinn *et al.*, 2020) | 296.5±21.4 | 89.3±0.6 | 70.6±2.9 | 77.6±2.9 | 64.7±7.4 |
| CLR-lasso (Susin *et al.*, 2020) | **2.0±0.2** | 100.0±0.0 | 77.5±1.8 | 81.6±2.2 | 75.8±2.7 |
| Random Forest | 10.6±0.4 | · | 78.0±2.2 | 82.2±2.2 | 77.3±2.5 |
| XGBoost (Chen and Guestrin, 2016) | 3.9±0.2 | · | 78.4±2.1 | 82.4±2.1 | 77.9±2.3 |

**Table 3:** Evaluation metrics for the liquid biopsy data (Best *et al.*, 2015), averaged over 20 independent 80/20 train/test splits. CoDaCoRe (with balances) achieves equal predictive accuracy as competing methods, but with much sparser solutions. Note that sparsity is expressed as an (integer) number of active variables in the model (not as a percentage of the total, as was done in Table 1).

|  | Runtime (s) | Vars (#) | Acc. (%) | AUC (%) | F1 (%) |
| --- | --- | --- | --- | --- | --- |
| Baseline | 0±0 | 0±0 | 79.1±0.0 | 50.0±0.0 | . |
| CoDaCoRe | 31±2.2 | **3±1** | **91.0±1.9** | 93.6±2.6 | **94.4±1.2** |
| Lasso | 23±0.2 | 22±4 | 87.8±1.3 | 94.7±1.5 | 92.7±0.7 |
| RF | 383±8.6 | . | 89.0±1.6 | 94.1±1.8 | 93.1±1.0 |
| XGBoost | 108±1.6 | . | 90.6±1.9 | **95.9±1.5** | 94.1±1.1 |

Figure 1 shows the average runtime of our classifiers on each dataset, with larger points denoting larger datasets. On these common benchmark datasets, CoDaCoRe trains up to 5 orders of magnitude faster than existing interpretable CoDa methods. On our larger datasets (3,090 inputs), selbal runs in ~ 100 hours, pairwise log-ratios and amalgam both run in ~10 hours, and CoDaCoRe runs in under 10 seconds (full runtimes are provided in Table 3 in the Supplement). All runs, including those involving gradient descent, were performed on identical CPU cores; CoDaCoRe can be accelerated further using GPUs, but we did not find it necessary to do so. It is also worth noting that the outperformance of CoDaCoRe is not merely as a result of the other methods failing on high-dimensional datasets. The consistent performance of CoDaCoRe across smaller and larger datasets is demonstrated in Supplementary Tables 4, 5, and 7, which show a full breakdown of results across each dataset.

**Figure 1:**
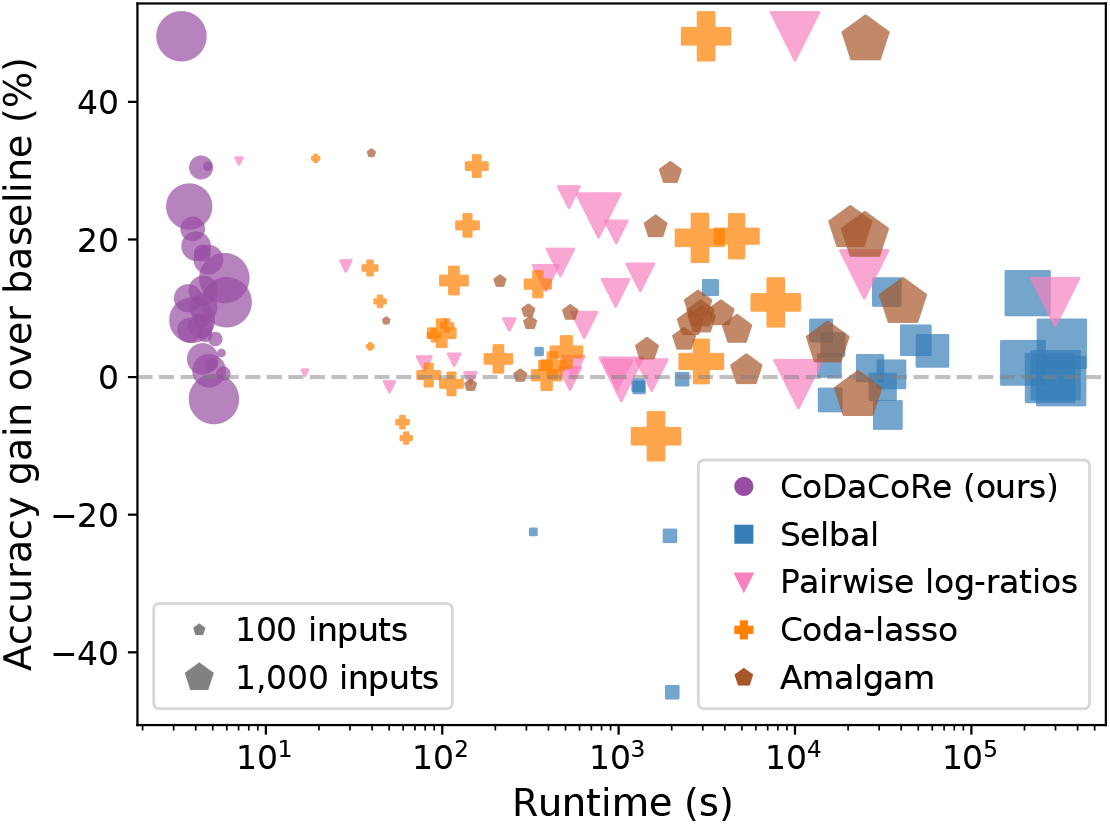
Classification accuracy (relative to the “majority vote” baseline) against runtime. Each point represents one of 25 datasets, with size proportional to the input dimension. Note the x-axis is drawn on the log-scale. CoDaCoRe (with balances) is the only method that scales effectively to our larger datasets, while consistently achieving high predictive accuracy. Moreover, its performance is broadly consistent across smaller and larger datasets.

Not only is CoDaCoRe sparser and more accurate than other interpretable models, it also performs on par with state-of-the-art black-box classifiers. By simply reducing the regularization parameter, from λ = 1 to λ = 0, CoDaCoRe (with balances) achieved an average 77.6% out-of-sample accuracy of and 82.0% AUC, on par with Random Forest and XGBoost (bottom rows of Table 2), while only using 5.9% of the input variables, on average. This result indicates, first, that CoDaCoRe provides a highly effective algorithm for variable selection in high-dimensional HTS data. Second, the fact that CoDaCoRe achieves similar predictive accuracy as state-of-the-art black-box classifiers, suggests that our model may have captured a near-complete representation of the signal in the data. At any rate, we take this as evidence that log-ratio transformed features are indeed of biological importance in the context of HTS data, corroborating previous microbiome research (Rahat-Rozenbloom *et al.*, 2014; Crovesy *et al.*, 2020; Magne *et al.*, 2020).

### 4.2 Interpretability

The CoDaCoRe algorithm offers two kinds of interpretability. First, it provides the analyst with sets of input variables whose aggregated ratio predicts the outcome of interest. These sets are easy to understand because they are discrete, with each component making an equivalent (unweighted) contribution. They are also sparse, usually containing fewer than 10 features per ratio, and can be made sparser by adjusting the regularization parameter λ. Such ratios have a precedent in microbiome research, for example the Firmicutes-to-Bacteroidetes ratio is used as a biomarker of gut health (Crovesy *et al.*, 2020; Magne *et al.*, 2020). Second, CoDaCoRe ranks predictive ratios hierarchically. Due to the ensembling procedure, the first ratio learned is the most predictive, the second ratio predicts the residual from the first, and so forth. Like principal components, the balances (or amalgamations) learned by CoDaCoRe are naturally ordered in terms of their explanatory power. This ordering aids interpretability by decomposing a multivariable model into comprehensible “chunks” of information.

Notably, we find a high degree of stability in the log-ratios selected by the model. We repeated CoDaCoRe on 10 independent training set splits of the Crohn disease data provided by Rivera-Pinto *et al.* (2018), and found consensus among the learned models. Figure 2 shows which bacteria were included for each split, in both versions of CoDaCoRe (balances and amalgamations). Importantly, most of the bacteria that were selected consistently by CoDaCoRe – notably Dialister, Roseburia and Clostridiales – were also identified by Rivera-Pinto *et al.* (2018). Differences between the sets selected by CoDaCoRe with balances vs. CoDaCoRe with amalgamations can be explained by differences in how the geometric mean vs. summation operations impact the log-ratio. The geometric mean, being more sensitive to small numbers, is more affected by the presence of rarer bacteria species like Dialister and Roseburia (as compared with the more common bacteria species like Haemophilus and Faecalibacterium).

**Figure 2:**
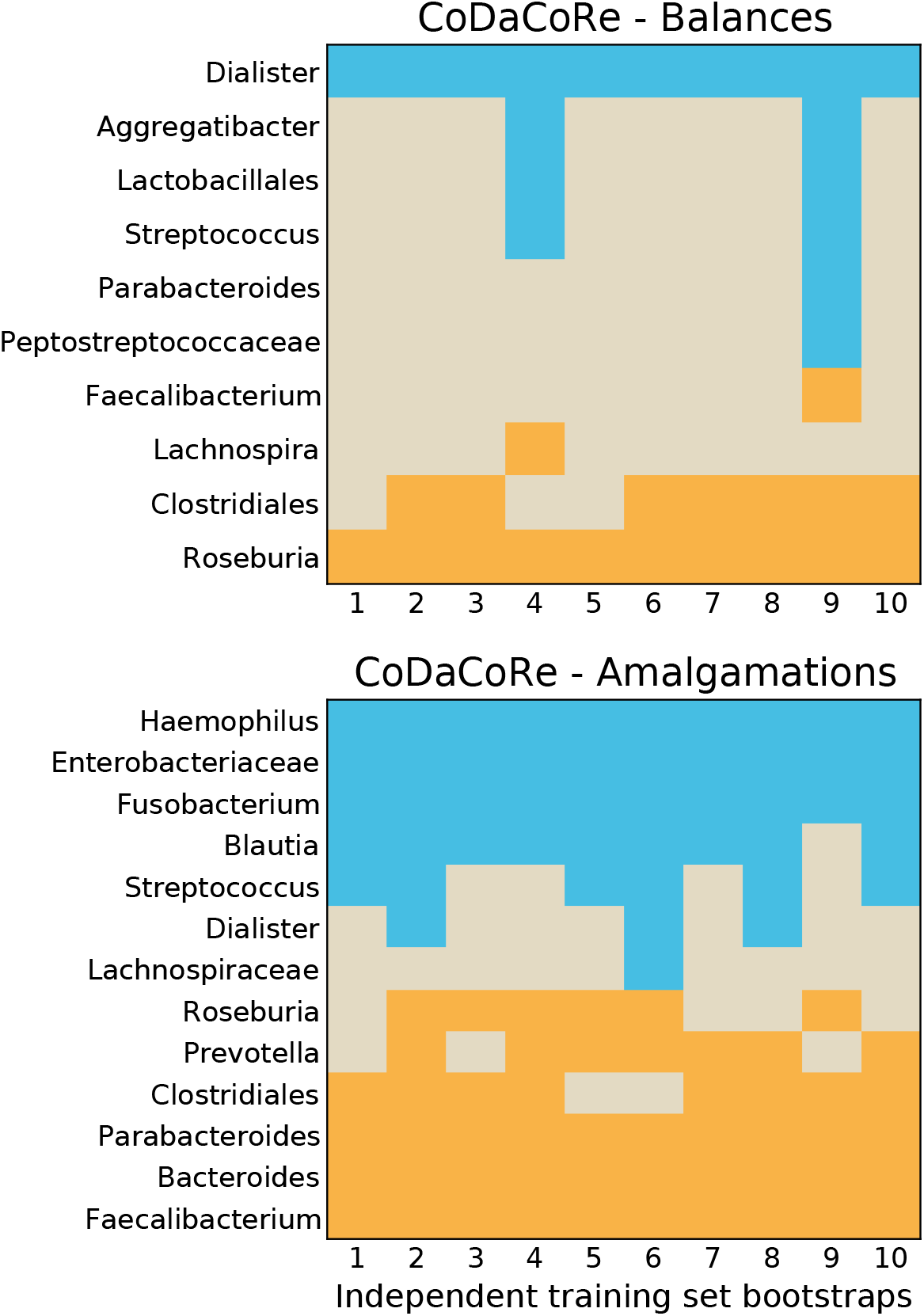
CoDaCoRe variable selection for the first (most explanatory) log-ratio on the Crohn disease data (Rivera-Pinto *et al.*, 2018). For each of 10 independent bootstraps of the training set (80% of the data randomly sampled with stratification by case-control), we show which variables are selected in the numerator (blue) and denominator (orange) of the log-ratio. Both versions of CoDaCoRe, with balances (top) or amalgamations (bottom), learn remarkably consistent log-ratios across independent training sets.

### 4.3 Scaling to Liquid Biopsy Data

HTS data generated from from clinical blood samples can be described as a “liquid biopsy” that can be used for cancer diagnosis and surveillance (Best *et al.*, 2015; Alix-Panabières and Pantel, 2016). These data can be very high-dimensional, especially when they include all gene transcripts as input variables. In a clinical context, the use of log-ratio predictors is an attractive option because they automatically correct for inter-sample sequencing biases that might otherwise limit the generalizability of the models (Dillies *et al.*, 2013). Unfortunately, existing log-ratio methods like selbal and amalgam simply cannot scale to liquid biopsy data sets that contain as many as 50,000 or more input variables.

The large dimensionality of such data has restricted its analysis to overly simplistic linear models, black-box models that are scalable but not interpretable, or suboptimal hybrid approaches where input variables must be pre-selected based on univariate measures (Best *et al.*, 2015; Zhang *et al.*, 2017; Sheng *et al.*, 2018). Owing to its linear scaling, CoDaCoRe can be fitted to these data at a similar computational cost to a single lasso regression, i.e., under a minute on a single CPU core. Thus, CoDaCoRe can be used to discover interpretable and predictive log-ratios that are suitable for liquid biopsy cancer diagnostics, among other similar applications.

We showcase the capabilities of CoDaCoRe in this high-dimensional setting, by applying our algorithm to the liquid biopsy data of (Best *et al.*, 2015). These data contain *p* = 58,037 genes sequenced in *n* = 288 human subjects, 60 of whom were healthy controls, the others having been previously diagnosed with cancer. Averaging over 20 random 80/20 train/test splits of this dataset, we found that CoDaCoRe achieved the same predictive accuracy as competing methods (within error), but obtained a much sparser model. Remarkably, CoDaCoRe identified log-ratios involving just 3 genes, that were equally predictive to both black-box classifiers and linear models with over 20 active variables. This case study again illustrates the potential of CoDaCoRe to derive novel biological insights, and also to develop learning algorithms for cancer diagnosis, a domain in which model interpretability – including sparsity - is of paramount importance (Wan *et al.*, 2017).

## 5 Conclusion

Our results corroborate the summary in Table 1: CoDaCoRe is the first sparse and interpretable CoDa model that can scale to high-dimensional HTS data. It does so convincingly, with linear scaling that results in runtimes similar to linear models. Our method is also competitive in terms of predictive accuracy, performing comparably to powerful black-box classifiers, but with interpretability. Our findings suggest that CoDaCoRe could play a significant role in the future analysis of high-throughput sequencing data, with broad implications in microbiology, statistical genetics, and the field of CoDa.

## Footnotes

1 The CoDaCoRe package is available at https://github.com/egr95/R-codacore.

2 Note that the original definition of balances includes a “normalization” constant, which we omit for clarity. This constant is in fact unnecessary, as it will get absorbed into a regression coefficient downstream.

3 Implemented using a heuristic search for improved computational efficiency (Quinn *et al.*, 2017).

## References

Aitchison, J. (1982). The statistical analysis of compositional data. Journal of the Royal Statistical Society: Series B (Methodological), 44(2), 139–160.

Alix-Panabières, C. and Pantel, K. (2016). Clinical applications of circulating tumor cells and circulating tumor dna as liquid biopsy. Cancer discovery, 6(5), 479–491.

Bates, S. and Tibshirani, R. (2019). Log-ratio lasso: Scalable, sparse estimation for log-ratio models. Biometrics, 75(2), 613–624.

Best, M. G., Sol, N., Kooi, I., Tannous, J., Westerman, B. A., Rustenburg, F., Schellen, P., Verschueren, H., Post, E., Koster, J., et al. (2015). Rna-seq of tumor-educated platelets enables blood-based pan-cancer, multiclass, and molecular pathway cancer diagnostics. Cancer cell, 28(5), 666–676.

Bondell, H. D. and Reich, B. J. (2009). Simultaneous factor selection and collapsing levels in anova. Biometrics, 65(1), 169–177.

Calle, M. L. (2019). Statistical analysis of metagenomics data. Genomics & informatics, 17(1).

Cammarota, G., Ianiro, G., Ahern, A., Carbone, C., Temko, A., Claesson, M. J., Gasbarrini, A., and Tortora, G. (2020). Gut microbiome, big data and machine learning to promote precision medicine for cancer. Nature Reviews Gastroenterology & Hepatology, 17(10), 635–648.

Chen, T. and Guestrin, C. (2016). Xgboost: A scalable tree boosting system. In Proceedings of the 22nd acm sigkdd international conference on knowledge discovery and data mining, pages 785–794.

Crovesy, L., Masterson, D., and Rosado, E. L. (2020). Profile of the gut microbiota of adults with obesity: a systematic review. European journal of clinical nutrition, 74(9), 1251–1262.

Dillies, M.-A., Rau, A., Aubert, J., Hennequet-Antier, C., Jeanmougin, M., Servant, N., Keime, C., Marot, G., Castel, D., Estelle, J., et al. (2013). A comprehensive evaluation of normalization methods for illumina high-throughput rna sequencing data analysis. Briefings in bioinformatics, 14(6), 671–683.

Egozcue, J. J. and Pawlowsky-Glahn, V. (2005). Groups of parts and their balances in compositional data analysis. Mathematical Geology, 37(7), 795–828.

Egozcue, J. J. and Pawlowsky-Glahn, V. (2016). Changing the reference measure in the simplex and its weighting effects. Austrian Journal of Statistics, 45(4), 25–44.

Egozcue, J. J. and Pawlowsky-Glahn, V. (2019). Compositional data: the sample space and its structure. TEST, 28(3), 599–638.

Egozcue, J. J., Pawlowsky-Glahn, V., Mateu-Figueras, G., and Barcelo-Vidal, C. (2003). Isometric logratio transformations for compositional data analysis. Mathematical Geology, 35(3), 279–300.

Fernandes, A. D., Macklaim, J. M., Linn, T. G., Reid, G., and Gloor, G. B. (2013). Anova-like differential expression (aldex) analysis for mixed population rna-seq. PLoS One, 8(7), e67019.

Fernandes, A. D., Reid, J. N., Macklaim, J. M., McMurrough, T. A., Edgell, D. R., and Gloor, G. B. (2014). Unifying the analysis of high-throughput sequencing datasets: characterizing rna-seq, 16s rrna gene sequencing and selective growth experiments by compositional data analysis. Microbiome, 2(1), 15.

Filzmoser, P. and Walczak, B. (2014). What can go wrong at the data normalization step for identification of biomarkers? Journal of Chromatography A, 1362, 194–205.

Filzmoser, P., Hron, K., and Reimann, C. (2009). Univariate statistical analysis of environmental (compositional) data: problems and possibilities. Science of the Total Environment, 407(23), 6100–6108.

Forslund, K., Hildebrand, F., Nielsen, T., Falony, G., Le Chatelier, E., Sunagawa, S., Prifti, E., Vieira-Silva, S., Gudmundsdottir, V., Pedersen, H. K., et al. (2015). Disentangling type 2 diabetes and metformin treatment signatures in the human gut microbiota. N atur e, 528(7581), 262–266.

Friedman, J., Hastie, T., Tibshirani, R., et al. (2001). The elements of statistical learning, volume 1. Springer series in statistics New York.

Gertheiss, J. and Tutz, G. (2010). Sparse modeling of categorial explanatory variables. The Annals of Applied Statistics, pages 2150–2180.

Gloor, G. B. and Reid, G. (2016). Compositional analysis: a valid approach to analyze microbiome high-throughput sequencing data. Canadian journal of microbiology, 62(8), 692–703.

Gloor, G. B., Wu, J. R., Pawlowsky-Glahn, V., and Egozcue, J. J. (2016). It’s all relative: analyzing microbiome data as compositions. Annals of epidemiology, 26(5), 322–329.

Gloor, G. B., Macklaim, J. M., Pawlowsky-Glahn, V., and Egozcue, J. J. (2017). Microbiome datasets are compositional: and this is not optional. Frontiers in microbiology, 8, 2224.

Goodman, B. and Flaxman, S. (2017). European union regulations on algorithmic decisionmaking and a “right to explanation”. AI magazine, 38(3), 50–57.

Gordon-Rodriguez, E., Loaiza-Ganem, G., and Cunningham, J. (2020a). The continuous categorical: a novel simplex-valued exponential family. In International Conference on Machine Learning, pages 3637–3647. PMLR.

Gordon-Rodriguez, E., Loaiza-Ganem, G., Pleiss, G., and Cunningham, J. P. (2020b). Uses and abuses of the cross-entropy loss: Case studies in modern deep learning. In Proceedings on “I Can’t Believe It’s Not Better!” at NeurIPS Workshops, volume 137 of Proceedings of Machine Learning Research, pages 1–10. PMLR.

Greenacre, M. (2019a). Comments on: Compositional data: the sample space and its structure. TEST, 28(3), 644–652.

Greenacre, M. (2019b). Variable selection in compositional data analysis using pairwise logratios. Mathematical Geosciences, 51(5), 649–682.

Greenacre, M. (2020). Amalgamations are valid in compositional data analysis, can be used in agglomerative clustering, and their logratios have an inverse transformation. Applied Computing and Geosciences, 5, 100017.

Greenacre, M., Grunsky, E., and Bacon-Shone, J. (2020). A comparison of isometric and amalgamation logratio balances in compositional data analysis. Computers & Geosciences, page 104621.

He, H. and Ma, Y. (2013). Imbalanced learning: foundations, algorithms, and applications.

Jang, E., Gu, S., and Poole, B. (2016). Categorical reparameterization with gumbel-softmax. arXiv preprint arXiv:1611.01144.

Li, H. (2015). Microbiome, metagenomics, and high-dimensional compositional data analysis. Annual Review of Statistics and Its Application, 2, 73–94.

Linderman, S., Mena, G., Cooper, H., Paninski, L., and Cunningham, J. (2018). Reparameterizing the birkhoff polytope for variational permutation inference. In International Conference on Artificial Intelligence and Statistics, pages 1618–1627. PMLR.

Lovell, D., Pawlowsky-Glahn, V., Egozcue, J. J., Marguerat, S., and Bähler, J. (2015). Proportionality: a valid alternative to correlation for relative data. PLoS Comput Biol, 11(3), e1004075.

Lu, J., Shi, P., and Li, H. (2019). Generalized linear models with linear constraints for microbiome compositional data. Biometrics, 75(1), 235–244.

Maddison, C. J., Mnih, A., and Teh, Y. W. (2017). The concrete distribution: A continuous relaxation of discrete random variables. In International Conference on Learning Representations.

Magne, F., Gotteland, M., Gauthier, L., Zazueta, A., Pesoa, S., Navarrete, P., and Balamurugan, R. (2020). The firmicutes/bacteroidetes ratio: a relevant marker of gut dysbiosis in obese patients? Nutrients, 12(5), 1474.

Martín-Fernández, J., Pawlowsky-Glahn, V., Egozcue, J., and Tolosona-Delgado, R. (2018). Advances in principal balances for compositional data. Mathematical Geosciences, 50(3), 273–298.

Martino, C., Morton, J. T., Marotz, C. A., Thompson, L. R., Tripathi, A., Knight, R., and Zengler, K. (2019). A novel sparse compositional technique reveals microbial perturbations. MSystems, 4(1).

Mena, G., Snoek, J., Linderman, S., and Belanger, D. (2018). Learning latent permutations with gumbel-sinkhorn networks. In International Conference on Learning Representations.

Mert, M. C., Filzmoser, P., and Hron, K. (2015). Sparse principal balances. Statistical Modelling, 15(2), 159–174.

Morton, J. T., Sanders, J., Quinn, R. A., McDonald, D., Gonzalez, A., Vázquez-Baeza, Y., Navas-Molina, J. A., Song, S. J., Metcalf, J. L., Hyde, E. R., et al. (2017). Balance trees reveal microbial niche differentiation. MSystems, 2(1).

Morton, J. T., Marotz, C., Washburne, A., Silverman, J., Zaramela, L. S., Edlund, A., Zengler, K., and Knight, R. (2019a). Establishing microbial composition measurement standards with reference frames. Nature communications, 10(1), 1–11.

Morton, J. T., Aksenov, A. A., Nothias, L. F., Foulds, J. R., Quinn, R. A., Badri, M. H., Swenson, T. L., Van Goethem, M. W., Northen, T. R., Vazquez-Baeza, Y., et al. (2019b). Learning representations of microbe–metabolite interactions. Nature methods, 16(12), 1306–1314.

Noguera-Julian, M., Rocafort, M., Guillén, Y., Rivera, J., Casadellà, M., Nowak, P., Hildebrand, F., Zeller, G., Parera, M., Bellido, R., et al. (2016). Gut microbiota linked to sexual preference and hiv infection. EBioMedicine, 5, 135–146.

Pawlowsky-Glahn, V. and Buccianti, A. (2011). Compositional data analysis: Theory and applications. John Wiley & Sons.

Pawlowsky-Glahn, V. and Egozcue, J. J. (2006). Compositional data and their analysis: an introduction. Geological Society, London, Special Publications, 264(1), 1–10.

Pawlowsky-Glahn, V., Egozcue, J. J., Tolosana Delgado, R., et al. (2011). Principal balances. Proceedings of CoDaWork, pages 1–10.

Pawlowsky-Glahn, V., Egozcue, J. J., and Tolosana-Delgado, R. (2015). Modeling and analysis of compositional data. John Wiley & Sons.

Pearson, K. (1896). Vii. mathematical contributions to the theory of evolution.—iii. regression, heredity, and panmixia. Philosophical Transactions of the Royal Society of London. Series A, containing papers of a mathematical or physical character, (187), 253–318.

Potapczynski, A., Loaiza-Ganem, G., and Cunningham, J. P. (2020). Invertible gaussian reparameterization: Revisiting the gumbel-softmax. Advances in Neural Information Processing Systems, 33.

Prifti, E., Chevaleyre, Y., Hanczar, B., Belda, E., Danchin, A., Clément, K., and Zucker, J.-D. (2020). Interpretable and accurate prediction models for metagenomics data. GigaScience, 9(3), giaa010.

Quinn, T., Nguyen, D., Rana, S., Gupta, S., and Venkatesh, S. (2020). Deepcoda: personalized interpretability for compositional health data. In International Conference on Machine Learning, pages 7877–7886. PMLR.

Quinn, T. P. and Erb, I. (2019). Using balances to engineer features for the classification of health biomarkers: a new approach to balance selection. bioRxiv, page 600122.

Quinn, T. P. and Erb, I. (2020). Amalgams: data-driven amalgamation for the dimensionality reduction of compositional data. NAR Genomics and Bioinformatics, 2(4), lqaa076.

Quinn, T. P., Richardson, M. F., Lovell, D., and Crowley, T. M. (2017). propr: an r-package for identifying proportionally abundant features using compositional data analysis. Scientific reports, 7(1), 1–9.

Quinn, T. P., Erb, I., Richardson, M. F., and Crowley, T. M. (2018). Understanding sequencing data as compositions: an outlook and review. Bioinformatics, 34(16), 2870–2878.

Quinn, T. P., Erb, I., Gloor, G., Notredame, C., Richardson, M. F., and Crowley, T. M. (2019). A field guide for the compositional analysis of any-omics data. GigaScience, 8(9), giz107.

Quinn, T. P., Gordon-Rodriguez, E., and Erb, I. (2021). A critique of differential abundance analysis, and advocacy for an alternative. arXiv preprint arXiv:2104.07266.

Rahat-Rozenbloom, S., Fernandes, J., Gloor, G. B., and Wolever, T. M. (2014). Evidence for greater production of colonic short-chain fatty acids in overweight than lean humans. International journal of obesity, 38(12), 1525–1531.

Rivera-Pinto, J., Egozcue, J. J., Pawlowsky-Glahn, V., Paredes, R., Noguera-Julian, M., and Calle, M. L. (2018). Balances: a new perspective for microbiome analysis. MSystems, 3(4).

Sheng, M., Dong, Z., and Xie, Y. (2018). Identification of tumor-educated platelet biomarkers of non-small-cell lung cancer. OncoTargets and therapy, 11, 8143.

Silverman, J. D., Washburne, A. D., Mukherjee, S., and David, L. A. (2017). A phylogenetic transform enhances analysis of compositional microbiota data. Elife, 6, e21887.

Susin, A., Wang, Y., Lé Cao, K.-A., and Calle, M. L. (2020). Variable selection in microbiome compositional data analysis. NAR Genomics and Bioinformatics, 2(2), lqaa029.

Templ, M. (2020). Artificial neural networks to impute rounded zeros in compositional data. arXiv preprint arXiv:2012.10300.

Tolosana-Delgado, R., Talebi, H., Khodadadzadeh, M., and Van den Boogaart, K. (2019). On machine learning algorithms and compositional data. In Proceedings of the 8th International Workshop on Compositional Data Analysis, Terrassa, Spain, pages 3–8.

Tutz, G. and Gertheiss, J. (2016). Regularized regression for categorical data. Statistical Modelling, 16(3), 161–200.

Van den Boogaart, K. G. and Tolosana-Delgado, R. (2013). Analyzing compositional data with R, volume 122. Springer.

Vangay, P., Hillmann, B. M., and Knights, D. (2019). Microbiome Learning Repo (ML Repo): A public repository of microbiome regression and classification tasks. GigaScience, 8(5).

Wan, J. C., Massie, C., Garcia-Corbacho, J., Mouliere, F., Brenton, J. D., Caldas, C., Pacey, S., Baird, R., and Rosenfeld, N. (2017). Liquid biopsies come of age: towards implementation of circulating tumour dna. Nature Reviews Cancer, 17(4), 223.

Washburne, A. D., Silverman, J. D., Leff, J. W., Bennett, D. J., Darcy, J. L., Mukherjee, S., Fierer, N., and David, L. A. (2017). Phylogenetic factorization of compositional data yields lineage-level associations in microbiome datasets. PeerJ, 5, e2969.

Zhang, Y.-H., Huang, T., Chen, L., Xu, Y., Hu, Y., Hu, L.-D., Cai, Y., and Kong, X. (2017). Identifying and analyzing different cancer subtypes using rna-seq data of blood platelets. Oncotarget, 8(50), 87494.

