## Supplement for "Learning Sparse Log-Ratios for High-Throughput Sequencing Data"

#### A Optimization

##### A.1 Continuous Relaxation

We optimized Eq. 10 from the main text using gradient descent with the following specifications:

- **Initialization.** We initialized the weights parameter $\mathbf{w}$ at zero, which corresponds to assigning no prior "preference" to one covariate over any another. As for the intercept parameter $\alpha$, initialization requires some care, particularly in order to account for possibly imbalanced classes and for the stage-wise additive nature of our model. Namely, $\alpha$ should captures the central tendency of the response so that the gradients propagated to $\mathbf{w}$ capture the *differential* effect of the covariates on the response (not doing so severely hinders gradient descent). In practice, this amounts to initializing $\alpha$ at the estimated coefficient of a generalized linear model with nothing but an intercept (which, in the regression case, reduces to the mean of the response). Note that, from the second stage of ensembling onwards, the fitted value of the current ensemble must be incorporated into said generalized linear model as a fixed offset term. Finally, we found the initialization of $\beta$ to matter less, and simply used the initial value $\beta = 0.1$ throughout.

- **Epochs.** We trained each continuous relaxation for 100 epochs. While this number of epochs was sufficient for the linear functionals we considered, we expect a higher number would be needed in order to combine our balances (or amalgamations) with nonlinear functionals.

- **Learning rate.** We used an adaptive scheme that exploits the structure of our continuous relaxation. Intuitively, we seek a learning rate that will allow $\mathbf{w}$ to explore the range of values where our relaxation has non-vanishing gradients, which approximately corresponds to $w_i \in (-5, 5)$. This interval corresponds to soft assignments approximately between $\tilde{w}_i \in (-0.99, 0.99)$; outside of this range, the sigmoid starts to saturate and gradients vanish. The question is then: how to choose a learning rate that allows $\mathbf{w}$ to explore the range $(-5, 5)$ efficiently (i.e., over a minimal number of epochs). We solve this indirectly; we pick the unique learning rate that would result in an initial gradient step that takes the maximum weight, $\max(|\mathbf{w}|)$, to the value 0.5. This learning rate can be found by simply computing one backward pass (and taking the maximum over the gradients w.r.t. $\mathbf{w}$) prior to applying any gradient steps. Note that the value of 0.5 is somewhat arbitrary, and simply represents the size of an initial step that should be "small, but significant relative to the target range of $(-5, 5)$".

- **Momentum.** We used momentum, with rate 0.9.

- **Minibatching.** We did not use minibatching, as we found it unnecessary on the datasets we considered. Nevertheless, minibatching could be beneficial, particularly for datasets with larger number of observations.

##### A.2 Discretization

Given a trained vector of soft assignments $\tilde{\mathbf{w}}$, typically only a small number will have converged to values close to $+1$ or $-1$, with most components remaining close to zero. As discussed in Section 3.3, in order to identify a balance it is sufficient to specify a threshold value $t$. However, different thresholds $t \in (0, 1)$ can result in different balances, depending on the values of $\tilde{\mathbf{w}}$. Our implementation evaluates a set of 20 "candidate thresholds", $t \in \{t_1, \ldots, t_{20}\}$, and settles on the one that yields the best cross-validation score. Any grid of values $\{t_1, \ldots, t_{20}\} \in (0, 1)$ can be used, where we write $\{t_1, \ldots, t_{20}\}$ in decreasing order. A principled choice is to "step through" the weights. Namely, start by rescaling the positive and negative components of $\tilde{\mathbf{w}}$ so that $\max \tilde{\mathbf{w}}^+ = \max \tilde{\mathbf{w}}^-$, and then set $\{t_1, \ldots, t_{20}\}$ to equal the top 20 order statistics of the set $\{|\tilde{w}_j|\}$. Thus, the first candidate $t_1$ will yield a simple log-ratio between two covariates (the rescaling ensure precisely two of our weights are $\geq t_1$), the second candidate $t_2$ will yield a log-ratio with a total of three covariates, and so forth.

Given a set of candidate thresholds $\{t_1, \ldots, t_{20}\}$, we associate a score to each of these via cross-validation. We first split the training data into 5 folds (sampled with stratification by case control (He and Ma, 2013)). Then, for each candidate threshold $t_i$, we fit 5 regressors of the form Eq. 3 on the 5 cross-validation training sets, we evaluate on their respective validation folds, and average the results to obtain the overall cross-validation score of $t_i$. Finally, we choose the largest threshold (i.e., the sparsest model) whose cross-validation score is within 1 standard error of the optimum achieved over $\{t_1, \ldots, t_{20}\}$ (with standard errors computed over the 5 cross-validation folds).

Note that we could search in this way over more than 20 candidate thresholds; the choice of the number 20 is by analogy to *selbal* (Rivera-Pinto *et al.*, 2018), where balances of up to (but no more than) 20 covariates are considered. We also found that our results were broadly insensitive to searching over more candidate thresholds; increasing this number can lead to slightly more accurate models that are slightly less sparse, and, evidently, somewhat slower to compute.

### B Datasets

Table 2 provides further details on the 25 datasets used in Section 4 of the the main text. Note also that zero-replacement is necessary prior to applying log-ratio transformations, a standard pre-processing step in the field of CoDa (Martín-Fernández *et al.*, 2000; Aitchison *et al.*, 2003; Martín-Fernández *et al.*, 2012). When necessary, we carried out zero-replacement by simply adding one unit to all counts prior to normalization.

### C Additional Experimental Results

Tables 3, 4, 5, 7 show our evaluation metrics on each individual dataset for a selection of our models. Table 2 from the main text summarizes these four tables by averaging over all datasets.

### References


Aitchison, J., Kay, J. W., *et al.* (2003). Possible solution of some essential zero problems in compositional data analysis.

He, H. and Ma, Y. (2013). Imbalanced learning: foundations, algorithms, and applications.

Martín-Fernández, J., Barceló-Vidal, C., and Pawlowsky-Glahn, V. (2000). Zero replacement in compositional data sets. In *Data analysis, classification, and related methods*, pages 155–160. Springer.

Martín-Fernández, J. A., Hron, K., Templ, M., Filzmoser, P., and Palarea-Albaladejo, J. (2012). Model-based replacement of rounded zeros in compositional data: classical and robust approaches. *Computational Statistics & Data Analysis*, **56**(9), 2688–2704.

Rivera-Pinto, J., Egozcue, J. J., Pawlowsky-Glahn, V., Paredes, R., Noguera-Julian, M., and Calle, M. L. (2018). Balances: a new perspective for microbiome analysis. *MSystems*, **3**(4).

Table 1: Data description. $n$ denotes the number of observations, $p$ the number of covariates. We also show the number of observations in the case and control groups.

| Dataset ID | $n$ | $p$ | Group 1 | Group 2 |
|---|---|---|---|---|
| 1 | 975 | 48 | Crohn's disease | Without |
| 2 | 128 | 60 | Men who have sex with men | Without |
| 3 | 220 | 153 | Control | IBD |
| 4 | 164 | 158 | Crohn's disease | Ulcerative colitis |
| 5 | 220 | 885 | Control | IBD |
| 6 | 164 | 885 | Crohn's disease | Ulcerative colitis |
| 7 | 182 | 278 | Case | Diarrheal control |
| 8 | 247 | 610 | Case | Non-Diarrheal control |
| 9 | 292 | 1133 | Colorectal cancer (CRC) | Without |
| 10 | 318 | 1302 | Colorectal cancer (CRC) | Non-CRC control |
| 11 | 1182 | 188 | Primary solid tumor | Solid tissue normal |
| 12 | 1004 | 188 | Her2 Cancer | Not Her2 Cancer |
| 13 | 718 | 188 | LumA Cancer | LumB Cancer |
| 14 | 140 | 992 | Crohn's disease (ileum) | Without (ileum) |
| 15 | 160 | 992 | Crohn's disease (rectum) | Without (rectum) |
| 16 | 2070 | 3090 | GI tract | Oral |
| 17 | 180 | 3090 | Female | Male |
| 18 | 404 | 3090 | Stool | Tongue (dorsum) |
| 19 | 408 | 3090 | Subgingival plaque | Supragingival plaque |
| 20 | 172 | 980 | Healthy | Colorectal cancer |
| 21 | 124 | 2526 | Without | Diabetes |
| 22 | 130 | 2579 | Cirrhosis | Without |
| 23 | 199 | 660 | Black | Hispanic |
| 24 | 342 | 660 | Nugent score high | Nugent score low |
| 25 | 200 | 660 | Black | White |

Table 2: Data description.

| Dataset ID | Group 1 Size | Group 2 Size | Source |
|---|---|---|---|
| 1 | 662 | 313 | doi: 10.1016/j.chom.2014.02.005 |
| 2 | 73 | 55 | doi:10.1016/j.ebiom.2016.01.032 |
| 3 | 56 | 164 | doi: 10.1038/s41564-018-0306-4 |
| 4 | 88 | 76 | doi: 10.1038/s41564-018-0306-4 |
| 5 | 56 | 164 | doi: 10.1038/s41564-018-0306-4 |
| 6 | 88 | 76 | doi: 10.1038/s41564-018-0306-4 |
| 7 | 93 | 89 | doi: 10.1128/mBio.01021-14 |
| 8 | 93 | 154 | doi: 10.1128/mBio.01021-15 |
| 9 | 120 | 172 | doi: 10.1186/s13073-016-0290-3 |
| 10 | 120 | 198 | doi: 10.1186/s13073-016-0290-3 |
| 11 | 1078 | 104 | doi: 10.1038/ng.2764 |
| 12 | 77 | 927 | doi: 10.1038/ng.2764 |
| 13 | 524 | 194 | doi: 10.1038/ng.2764 |
| 14 | 78 | 62 | doi: 10.1016/j.chom.2014.02.005 |
| 15 | 68 | 92 | doi: 10.1016/j.chom.2014.02.005 |
| 16 | 227 | 1843 | doi: 10.1038/nature11209 |
| 17 | 82 | 98 | doi: 10.1038/nature11209 |
| 18 | 204 | 200 | doi: 10.1038/nature11209 |
| 19 | 203 | 205 | doi: 10.1038/nature11209 |
| 20 | 86 | 86 | doi: 10.1101/gr.126573.111 |
| 21 | 59 | 65 | doi: 10.1038/nature11450 |
| 22 | 68 | 62 | doi: 10.1038/nature13568 |
| 23 | 104 | 95 | doi: 10.1073/pnas.1002611107 |
| 24 | 97 | 245 | doi: 10.1073/pnas.1002611107 |
| 25 | 104 | 96 | doi: 10.1073/pnas.1002611107 |

Table 3: Average runtime over 20 train/test splits, in seconds.

| Dataset ID | CoDaCoRe (Defaults) | selbal | Pairwise log-ratios | Coda-lasso | amalgam | Random Forest |
|---|---|---|---|---|---|---|
| 1 | 6 | 328 | 17 | 39 | 48 | 2 |
| 2 | 5 | 354 | 7 | 19 | 40 | 0 |
| 3 | 5 | 1307 | 50 | 62 | 145 | 1 |
| 4 | 4 | 1291 | 28 | 44 | 212 | 1 |
| 5 | 4 | 31407 | 638 | 420 | 2553 | 5 |
| 6 | 4 | 26638 | 387 | 348 | 3806 | 4 |
| 7 | 4 | 3316 | 79 | 39 | 532 | 1 |
| 8 | 4 | 14058 | 522 | 157 | 1965 | 3 |
| 9 | 4 | 48394 | 985 | 391 | 4694 | 11 |
| 10 | 5 | 60138 | 1540 | 505 | 5300 | 15 |
| 11 | 5 | 1954 | 240 | 90 | 315 | 5 |
| 12 | 6 | 2290 | 144 | 59 | 277 | 6 |
| 13 | 5 | 2014 | 117 | 107 | 307 | 5 |
| 14 | 4 | 33619 | 961 | 99 | 2919 | 2 |
| 15 | 5 | 35193 | 1326 | 116 | 2957 | 3 |
| 16 | 6 | 279072 | 298672 | 7776 | 40949 | 135 |
| 17 | 5 | 300431 | 10449 | 1628 | 22777 | 10 |
| 18 | 3 | 322726 | 9997 | 3131 | 25130 | 8 |
| 19 | 6 | 325642 | 24691 | 2890 | 24908 | 25 |
| 20 | 4 | 33150 | 468 | 208 | 2822 | 3 |
| 21 | 4 | 196216 | 1036 | 2942 | 15328 | 6 |
| 22 | 4 | 208464 | 769 | 4668 | 20612 | 6 |
| 23 | 4 | 15818 | 530 | 84 | 1447 | 2 |
| 24 | 4 | 16382 | 969 | 139 | 1621 | 4 |
| 25 | 4 | 15640 | 553 | 113 | 2349 | 2 |
| Mean | 5 | 79034 | 14207 | 1043 | 7361 | 11 |

Table 4: Proportion of input variables active (%), averaged over 20 train/test splits

| Dataset ID | CoDaCoRe (Defaults) | selbal | Pairwise log-ratios | Coda-lasso | amalgam | Random Forest |
|---|---|---|---|---|---|---|
| 1 | 15.4±2.4 | 25.6±3.1 | 21.0±2.4 | 53.8±5.5 | 85.2±2.8 | 100.0±0.0 |
| 2 | 4.5±0.6 | 5.5±0.4 | 11.2±1.9 | 9.4±0.7 | 86.1±2.2 | 100.0±0.0 |
| 3 | 2.1±0.3 | 3.0±0.5 | 3.5±0.9 | 6.7±1.6 | 80.7±2.7 | 100.0±0.0 |
| 4 | 2.0±0.3 | 1.8±0.2 | 4.2±0.8 | 6.3±1.4 | 81.6±1.6 | 100.0±0.0 |
| 5 | 0.6±0.3 | 0.2±0.0 | 0.7±0.1 | 1.9±0.7 | 84.9±0.9 | 100.0±0.0 |
| 6 | 0.2±0.0 | 0.3±0.0 | 0.8±0.1 | 0.7±0.1 | 85.3±0.8 | 100.0±0.0 |
| 7 | 3.3±0.6 | 0.8±0.1 | 0.2±0.1 | 2.2±0.5 | 79.6±2.5 | 100.0±0.0 |
| 8 | 1.3±0.2 | 1.0±0.1 | 1.7±0.2 | 4.2±1.0 | 81.9±2.2 | 100.0±0.0 |
| 9 | 0.8±0.1 | 0.2±0.0 | 0.2±0.1 | 1.2±0.4 | 89.7±1.0 | 100.0±0.0 |
| 10 | 0.6±0.1 | 0.2±0.0 | 0.0±0.0 | 1.1±0.2 | 90.0±1.3 | 100.0±0.0 |
| 11 | 1.8±0.9 | 4.2±0.5 | 2.5±0.3 | 74.2±10.6 | 91.0±1.7 | 100.0±0.0 |
| 12 | 5.3±0.9 | 6.9±0.5 | 5.0±0.9 | 62.1±8.8 | 89.0±1.5 | 100.0±0.0 |
| 13 | 5.2±0.4 | 7.1±0.4 | 5.9±0.8 | 58.1±12.7 | 84.5±1.7 | 100.0±0.0 |
| 14 | 0.8±0.2 | 0.2±0.0 | 0.6±0.1 | 0.4±0.1 | 92.9±1.4 | 100.0±0.0 |
| 15 | 0.6±0.2 | 0.3±0.1 | 0.6±0.1 | 0.6±0.2 | 90.8±1.1 | 100.0±0.0 |
| 16 | 0.1±0.0 | 0.4±0.0 | 0.2±0.0 | 95.1±9.9 | 94.8±1.3 | 100.0±0.0 |
| 17 | 0.1±0.0 | 0.1±0.0 | 0.0±0.0 | 0.0±0.0 | 94.0±2.6 | 100.0±0.0 |
| 18 | 0.1±0.0 | 0.3±0.0 | 0.3±0.0 | 100.0±0.0 | 88.6±4.5 | 100.0±0.0 |
| 19 | 0.1±0.0 | 0.1±0.0 | 0.3±0.0 | 0.4±0.1 | 97.8±1.1 | 100.0±0.0 |
| 20 | 0.5±0.1 | 0.2±0.0 | 0.4±0.1 | 0.4±0.5 | 89.1±2.5 | 100.0±0.0 |
| 21 | 0.3±0.1 | 0.1±0.0 | 0.0±0.0 | 1.1±0.9 | 92.1±3.7 | 100.0±0.0 |
| 22 | 0.1±0.0 | 0.1±0.0 | 0.3±0.1 | 10.0±10.7 | 89.9±1.8 | 100.0±0.0 |
| 23 | 0.5±0.1 | 0.3±0.0 | 0.2±0.1 | 0.5±0.3 | 86.7±3.3 | 100.0±0.0 |
| 24 | 1.0±0.1 | 0.8±0.1 | 1.9±0.1 | 1.6±0.1 | 93.2±1.2 | 100.0±0.0 |
| 25 | 0.3±0.0 | 0.3±0.0 | 0.7±0.1 | 0.2±0.1 | 69.8±5.7 | 100.0±0.0 |
| Mean | 1.9±0.3 | 2.4±0.2 | 2.5±0.4 | 19.7±2.7 | 87.6±2.1 | 100.0±0.0 |

Table 5: Out-of-sample accuracy (%) averaged over 20 train/test splits.

| Dataset ID | CoDaCoRe (Defaults) | selbal | Pairwise log-ratios | Coda-lasso | amalgam | Random Forest |
|---|---|---|---|---|---|---|
| 1 | 71.4±1.1 | 45.4±2.9 | 68.5±1.0 | 72.4±1.7 | 76.1±1.4 | 81.5±1.4 |
| 2 | 87.5±2.5 | 60.6±3.6 | 88.3±3.1 | 88.7±2.8 | 89.4±2.8 | 90.8±2.3 |
| 3 | 76.7±2.4 | 73.3±0.0 | 73.4±0.2 | 66.0±3.0 | 73.7±2.4 | 81.7±1.5 |
| 4 | 61.9±2.6 | 52.6±1.9 | 70.0±2.7 | 64.8±3.0 | 67.8±3.7 | 75.0±2.4 |
| 5 | 85.0±2.0 | 73.4±0.2 | 82.4±1.5 | 76.6±3.4 | 82.7±2.4 | 87.9±2.0 |
| 6 | 65.3±3.5 | 55.1±2.0 | 68.2±3.0 | 67.3±3.0 | 63.1±2.9 | 73.2±3.1 |
| 7 | 69.0±3.8 | 64.0±2.8 | 53.0±2.0 | 66.9±4.0 | 60.4±2.9 | 69.7±5.6 |
| 8 | 92.9±1.4 | 69.2±1.8 | 88.6±2.3 | 93.1±1.4 | 92.1±1.5 | 96.2±1.0 |
| 9 | 61.4±2.9 | 64.2±1.4 | 59.1±0.3 | 59.1±3.9 | 65.7±3.2 | 67.5±2.1 |
| 10 | 63.1±1.6 | 66.0±0.7 | 62.5±0.0 | 65.9±3.3 | 63.2±2.7 | 66.4±1.1 |
| 11 | 96.8±0.5 | 68.1±9.2 | 98.9±0.3 | 97.3±0.6 | 99.1±0.2 | 99.2±0.2 |
| 12 | 92.9±0.5 | 92.1±0.0 | 92.2±0.2 | 85.8±1.0 | 92.6±0.6 | 92.4±0.2 |
| 13 | 79.1±1.0 | 27.2±0.1 | 75.5±0.7 | 80.5±1.2 | 82.6±1.0 | 83.7±1.2 |
| 14 | 68.5±3.8 | 50.3±1.8 | 68.1±3.8 | 62.2±4.0 | 65.0±3.5 | 63.8±4.3 |
| 15 | 74.5±4.9 | 57.9±1.8 | 72.0±3.3 | 71.5±4.8 | 65.8±3.2 | 70.8±5.0 |
| 16 | 99.9±0.1 | 88.9±0.0 | 100.0±0.0 | 99.9±0.1 | 99.9±0.1 | 99.9±0.1 |
| 17 | 51.4±3.3 | 54.8±1.7 | 53.5±0.7 | 46.0±0.0 | 51.9±4.0 | 61.2±2.9 |
| 18 | 100.0±0.0 | 49.9±0.5 | 100.0±0.0 | 100.0±0.0 | 99.5±0.5 | 99.8±0.4 |
| 19 | 64.7±2.5 | 55.2±1.5 | 65.2±2.0 | 70.5±1.8 | 70.8±1.6 | 73.2±2.1 |
| 20 | 69.0±4.1 | 62.4±2.2 | 66.7±3.6 | 52.6±2.4 | 60.6±3.6 | 68.3±3.0 |
| 21 | 60.8±4.1 | 54.6±2.6 | 52.2±0.4 | 54.8±4.9 | 57.4±6.1 | 54.4±5.4 |
| 22 | 77.2±3.2 | 64.6±4.0 | 75.9±5.5 | 72.9±3.9 | 74.1±3.6 | 88.0±2.1 |
| 23 | 59.0±3.4 | 48.9±1.5 | 52.0±1.8 | 52.5±1.3 | 56.2±3.0 | 54.0±3.0 |
| 24 | 93.3±1.4 | 76.5±1.6 | 92.9±1.3 | 93.8±1.3 | 93.6±0.8 | 93.9±1.1 |
| 25 | 59.6±3.1 | 53.9±1.7 | 53.7±2.9 | 51.2±0.0 | 57.7±3.7 | 56.5±2.4 |
| Mean | 75.2±2.4 | 61.2±1.9 | 73.3±1.7 | 72.5±2.3 | 74.4±2.5 | 78.0±2.2 |

Table 6: Out-of-sample AUC (%) averaged over 20 train/test splits.

| Dataset ID | CoDaCoRe (Defaults) | selbal | Pairwise log-ratios | Coda-lasso | amalgam | Random Forest |
|---|---|---|---|---|---|---|
| 1 | 75.8±1.7 | 81.5±1.5 | 69.9±2.5 | 81.2±1.8 | 81.0±1.5 | 87.8±1.2 |
| 2 | 93.9±1.9 | 92.3±2.1 | 94.2±2.0 | 94.8±1.7 | 94.7±2.1 | 96.2±2.0 |
| 3 | 79.2±2.9 | 76.0±3.2 | 77.3±3.2 | 73.9±5.5 | 76.9±3.9 | 88.1±1.6 |
| 4 | 69.4±3.3 | 73.4±4.5 | 74.3±3.0 | 74.0±3.3 | 72.6±3.4 | 83.2±2.8 |
| 5 | 91.2±1.6 | 93.1±1.1 | 90.0±1.8 | 92.1±1.8 | 88.8±2.8 | 93.7±1.3 |
| 6 | 72.0±3.7 | 74.9±4.1 | 76.2±3.7 | 76.0±3.7 | 68.0±3.5 | 81.8±3.3 |
| 7 | 73.7±4.0 | 82.1±3.0 | 53.0±2.9 | 75.6±4.9 | 65.8±3.5 | 75.3±4.7 |
| 8 | 98.2±0.7 | 96.5±0.9 | 94.8±2.0 | 98.8±0.5 | 97.6±0.8 | 99.3±0.3 |
| 9 | 64.6±3.0 | 61.6±3.5 | 51.5±1.6 | 61.9±4.3 | 67.7±3.5 | 70.5±2.1 |
| 10 | 66.2±3.7 | 62.8±2.0 | 50.0±0.0 | 68.5±2.7 | 64.3±3.1 | 66.0±2.0 |
| 11 | 98.5±0.5 | 99.6±0.3 | 99.5±0.4 | 99.4±0.6 | 99.4±0.5 | 99.3±0.7 |
| 12 | 89.2±1.6 | 91.1±1.5 | 88.1±1.6 | 94.4±1.2 | 90.0±1.6 | 90.1±1.3 |
| 13 | 82.5±1.4 | 90.6±1.2 | 78.0±1.7 | 88.4±1.2 | 87.1±1.9 | 89.7±1.2 |
| 14 | 73.5±4.3 | 68.8±4.7 | 70.8±5.1 | 63.4±5.8 | 68.8±4.7 | 71.3±5.1 |
| 15 | 79.9±5.5 | 79.2±3.7 | 79.0±3.5 | 75.4±6.2 | 69.0±3.6 | 78.4±4.6 |
| 16 | 100.0±0.0 | 100.0±0.0 | 100.0±0.0 | 100.0±0.0 | 100.0±0.0 | 100.0±0.0 |
| 17 | 57.2±2.5 | 54.2±3.5 | 50.6±1.7 | 50.0±0.0 | 56.1±4.0 | 65.1±2.8 |
| 18 | 100.0±0.0 | 100.0±0.0 | 100.0±0.0 | 100.0±0.0 | 99.8±0.3 | 100.0±0.0 |
| 19 | 71.6±3.6 | 80.0±1.6 | 70.2±2.4 | 79.3±1.6 | 76.2±1.6 | 79.8±2.1 |
| 20 | 75.0±4.0 | 76.9±3.0 | 70.2±4.8 | 53.7±3.4 | 65.1±3.0 | 72.9±3.0 |
| 21 | 67.7±4.9 | 62.2±3.8 | 50.6±1.2 | 61.2±5.8 | 67.2±5.4 | 61.3±4.6 |
| 22 | 85.8±2.6 | 85.0±2.4 | 82.0±6.4 | 89.2±2.1 | 81.8±3.8 | 91.8±2.3 |
| 23 | 64.6±3.6 | 57.3±3.2 | 52.3±2.8 | 52.6±2.4 | 61.7±3.4 | 57.3±3.1 |
| 24 | 96.2±1.3 | 97.2±0.8 | 96.0±1.1 | 97.5±0.7 | 96.4±1.5 | 98.1±0.6 |
| 25 | 62.7±3.8 | 63.5±3.2 | 60.8±3.4 | 49.9±0.1 | 59.4±3.6 | 58.2±2.6 |
| Mean | 79.5±2.6 | 80.0±2.4 | 75.2±2.4 | 78.0±2.4 | 78.2±2.7 | 82.2±2.2 |

Table 7: Out-of-sample F1 score (%) averaged over 20 train/test splits.

| Dataset ID | CoDaCoRe (Defaults) | selbal | Pairwise log-ratios | Coda-lasso | amalgam | Random Forest |
|---|---|---|---|---|---|---|
| 1 | 46.9±2.8 | 53.8±1.2 | 10.9±6.5 | 65.3±1.8 | 58.6±2.6 | 65.7±2.8 |
| 2 | 85.6±2.6 | 68.0±2.1 | 86.0±3.6 | 86.4±3.8 | 87.1±3.6 | 89.1±2.8 |
| 3 | 84.1±1.8 | 84.6±0.0 | 84.6±0.1 | 73.3±2.9 | 82.2±1.6 | 88.4±0.9 |
| 4 | 56.8±3.6 | 65.7±1.1 | 65.0±3.1 | 65.0±3.6 | 66.4±4.0 | 71.7±3.0 |
| 5 | 89.6±1.4 | 84.7±0.1 | 88.7±0.9 | 81.0±3.5 | 87.9±1.7 | 91.8±1.4 |
| 6 | 60.9±4.5 | 66.2±1.6 | 63.6±4.6 | 66.2±3.5 | 56.9±4.5 | 70.4±3.9 |
| 7 | 69.0±3.9 | 72.9±1.7 | 10.6±9.5 | 70.0±7.8 | 60.3±3.6 | 69.3±5.2 |
| 8 | 94.3±1.2 | 80.2±0.9 | 91.2±1.7 | 94.6±1.0 | 93.7±1.2 | 97.0±0.8 |
| 9 | 68.3±2.5 | 76.8±0.7 | 74.2±0.3 | 62.2±10.2 | 70.9±2.9 | 76.6±1.6 |
| 10 | 72.1±1.3 | 78.6±0.4 | 76.9±0.0 | 73.6±4.0 | 70.6±2.4 | 77.9±0.7 |
| 11 | 80.8±2.8 | 38.9±3.9 | 93.5±1.8 | 87.0±2.5 | 94.9±1.3 | 95.2±1.4 |
| 12 | 96.2±0.3 | 95.9±0.0 | 95.9±0.1 | 91.8±0.7 | 96.1±0.3 | 96.0±0.1 |
| 13 | 54.7±2.1 | 42.7±0.1 | 31.6±3.9 | 68.2±1.9 | 64.5±2.8 | 62.7±3.5 |
| 14 | 63.2±3.9 | 64.1±1.1 | 52.9±9.8 | 38.1±13.7 | 60.5±4.3 | 53.0±6.9 |
| 15 | 77.5±4.4 | 72.5±1.2 | 75.9±3.0 | 75.0±4.7 | 69.8±3.5 | 75.9±4.0 |
| 16 | 100.0±0.0 | 94.1±0.0 | 100.0±0.0 | 100.0±0.0 | 100.0±0.0 | 99.9±0.1 |
| 17 | 58.4±3.6 | 69.8±1.2 | 69.4±1.1 | 0.0±0.0 | 57.0±4.2 | 65.8±2.9 |
| 18 | 100.0±0.0 | 66.3±0.2 | 100.0±0.0 | 100.0±0.0 | 99.5±0.5 | 99.8±0.4 |
| 19 | 64.5±2.5 | 69.0±0.7 | 65.7±2.1 | 73.0±1.8 | 71.3±1.7 | 73.4±2.3 |
| 20 | 69.5±3.6 | 71.9±1.2 | 68.9±2.6 | 14.8±11.9 | 60.5±3.6 | 66.9±3.1 |
| 21 | 62.1±4.5 | 67.1±2.5 | 68.2±0.4 | 34.1±14.8 | 59.3±5.9 | 55.8±6.1 |
| 22 | 76.8±3.1 | 72.7±2.2 | 74.4±6.4 | 77.4±2.6 | 73.2±4.1 | 88.1±1.9 |
| 23 | 60.8±5.0 | 64.1±1.0 | 7.3±4.8 | 10.2±7.8 | 51.6±3.6 | 50.4±3.3 |
| 24 | 95.4±0.9 | 85.8±0.8 | 95.2±0.9 | 95.8±0.8 | 95.6±0.6 | 95.7±0.8 |
| 25 | 54.7±3.8 | 66.8±1.0 | 43.0±8.0 | 2.5±5.0 | 59.7±4.2 | 55.5±2.9 |
| Mean | 73.7±2.6 | 70.9±1.1 | 67.8±3.0 | 64.2±4.4 | 73.9±2.8 | 77.3±2.5 |
